# The desmosomal cadherin Desmoglein-3 regulates YAP and phospho-YAP in keratinocyte responses to mechanical forces

**DOI:** 10.1101/827725

**Authors:** Jutamas Uttagomol, Usama Sharif Ahmad, Ambreen Rehman, Yunying Huang, Ana C. Laly, Angray Kang, Jan Soetaert, Randy Chance, Muy-Teck Teh, John T. Connelly, Hong Wan

## Abstract

Desmoglein 3 (Dsg3), plays a crucial role in cell-cell adhesion and tissue integrity. Increasing evidence suggests that Dsg3 acts as a regulator of cellular mechanotransduction, but little is known about its direct role in mechanical force transmission. The present study investigated the impact of cyclic strain and substrate stiffness on Dsg3 expression and its role in mechanotransduction. A direct comparison was made with E-cadherin, a well-characterized mechanosensor, in human keratinocytes. Exposure of oral and skin keratinocytes to equiaxial cyclic strain promoted changes in expression and localization of junction assembly proteins. Knockdown of Dsg3 by siRNA blocked strain-induced junctional remodeling of E-cadherin and Myosin IIa. Importantly, the study demonstrated that Dsg3 regulates the expression and localization of YAP, a mechanosensor and an effector of the Hippo pathway. Furthermore, we showed that Dsg3 forms a complex with phospho-YAP and sequestered it to the plasma membrane, while Dsg3 depletion had an impact on both YAP and phospho-YAP in their response to mechanical forces, increasing the sensitivity of keratinocytes to both strain-or substrate rigidity-induced nuclear relocalization of YAP and phospho-YAP. We showed that PKP1 seemed to be the key in such a complex formation since its silencing resulted in Dsg3 disruption at the junctions with concomitant loss of pYAP in the peripheral cytoplasm. Finally, we demonstrated that this Dsg3/YAP pathway has an influence on the expression of *YAP1* target genes as well as cell proliferation in keratinocytes. Together, these findings provide evidence of a novel role for Dsg3 in keratinocyte mechanotransduction.

## 1. Introduction

The hallmark of epithelia is that individual cells attach to each another through numerous intercellular junctions with two anchoring junctions, *i.e*. adherens junctions (AJs) and desmosomes (DSMs), being especially important for cellular architecture and tissue integrity. The cell-cell adhesions in both junctions are mediated by cadherin family proteins in a calcium-dependent manner. The adhesion receptors in AJs belong to the classical cadherin family, such as E-cadherin, whereas those in DSMs comprise of two desmosomal cadherin subfamilies, desmoglein (Dsg) and desmocollin (Dsc). All these cadherins are the single-pass transmembrane proteins and are linked to cytoskeletal filaments on their cytoplasmic tails through plaque proteins, including armadillo proteins (α, β, γ-catenins, and plakophilins) and plakin family members (*e.g*. desmoplakin). Both AJs and DSMs are enriched in stratified epithelial tissues, such as skin and mucous membrane that are subjected to mechanical stresses on a daily basis.

Cadherin-mediated cell adhesion plays an essential role in contact inhibition of cell proliferation (CIP) via the Hippo signaling pathway; the latter being an essential regulator in control of cell differentiation and organ growth, with its deregulation contributing to cancer development [2,39]. Depletion of the components in the Hippo pathway, such as β-and α-catenins, inhibits the E-cadherin mediated CIP, leading to increased cell proliferation [27]. This is due to the activation of Yes-associated protein (YAP), a transcriptional coactivator and a key downstream effector of the Hippo pathway with its activation causing YAP phosphorylation at specific serine residues, such as S127, resulting in nuclear exclusion [27]. YAP also has been identified as a mechanosensor independent of the Hippo cascade and such a function is driven by Rho GTPase activity and the tension generated by the actomyosin cytoskeleton [16,33]. The external physical forces, such as mechanical loading and substrate stiffness, can activate YAP and drive its nuclear translocation, leading to transcription of YAP target genes responsible for cell proliferation [33,37]. It is increasingly appreciated that mechanotransduction is as important as chemical factors in controlling diverse cell behavior, including growth, differentiation and tumor progression [29,47].

E-cadherin mediated AJs, in concert with associated F-actin and actomyosin, serve as important hubs in mechanosensing and mechanotransduction that influence cell-cell adhesion and traction force at cell-ECM junctions [14,26,29-31,57]. However, little is known about the role of desmosomal cadherins in these processes in spite of the fact that DSMs are crucial in cell signaling and tissue integrity [52]. The importance of desmosomal cadherins, as well as other DSM constitutive proteins, in tissue homeostasis and physiological function, has been underscored by various vesiculobullous diseases with clinical manifestation of blistering and fragility syndrome of skin and oral mucosa as well as heart failure in some cases [10,52]. Recent studies have uncovered that two DSM proteins, Dsg2 and Dp, are capable of sensing external mechanical loading, highlighting the probable involvement of DSM proteins in mechanobiology [3,40].

Desmoglein-3 (Dsg3) is an isoform of the Dsg subfamily with uniform expression across the entire stratified epithelium in oral mucosa and restricted expression in the basal layer of the epidermis in the skin [51]. Why Dsg3 exhibits such distinct tissue expression patterns remains unclear. Perhaps, Dsg3 is best known as PVA, the autoantigen in pemphigus vulgaris, a life-threatening autoimmune blistering disease. The importance of Dsg3 in cell-cell adhesion is highlighted by numerous studies based on pemphigus autoantibodies that demonstrate convincingly that disruption of Dsg3 is a causative factor for the skin and oral lesions [1,48]. However, the action of Dsg3 is not restricted to DSM adhesion and in fact, Dsg3 is present at the plasma membrane beyond the DSMs [1,6,43,54,55]. *In vitro* studies have shown that extra-junctional Dsg3 “cross-talks” with E-cadherin and regulates various signaling pathways, e.g. Src, Ezrin and Rho GTPases Rac1/cdc42 and transcription factor c-Jun/AP-1, all of which are involved in modulating the actin cytoskeleton; implying a role for Dsg3 in mechanotransduction [7,53-55]. Moreover, the interaction of actin proteins with the cytoplasmic domain of Dsg3 has been demonstrated by Mass Spec analysis [7]. Overexpression of Dsg3 results in enhanced membrane projections and cell spreading, with a consequence of accelerated cell migration and invasion in cancer cell lines [7,55]. Upregulation of Dsg3 also has been found in solid cancer where tissue stiffness has been altered [11,19,23,46,49]. Collectively, these lines of evidence led us to hypothesize that Dsg3 may be involved in cellular force transmission and transduction, somewhat in analogous to E-cadherin. In this present study, we have addressed this hypothesis in keratinocytes and found that Dsg3 responds to mechanical loading and is required for AJ remodeling induced by cyclic strain or substrate stiffness. Importantly, we have identified Dsg3 as a regulator of YAP and phospho-YAP (pYAP) in cell proliferation and in response to mechanical stress via, at least one, mechanism of forming a complex with pYAP and regulating its cellular localization.

## 2. Materials and Methods

### 2.1. Antibodies

The following mouse and rabbit monoclonal/polyclonal antibodies (Abs) were used: 5H10, mouse Ab against N-terminus of Dsg3 (sc-23912, Santa Cruz); H-145, rabbit Ab against C-terminus of Dsg3 (sc-20116, Santa Cruz); 33–3D, mouse IgM against Dsg2 (gift from Professor Garrod); rabbit Ab to Dsc2 (610120, Progen); Dsc3-U114, mouse Ab to Dsc3 (65193, Progen); PG 5.1, mouse Ab to Plakoglobin (65015, Progen); 115F, mouse Ab to Desmoplakin (gift from Professor Garrod); H-300, rabbit Ab to Desmoplakin (sc-33555, Santa Cruz); 5C2, mouse Ab to plakophilin1 (Progen); PKP3, mouse Ab (ab151401, Abcam), HECD-1, mouse anti-N-terminus of E-cadherin (ab1416, Abcam); 6F9, mouse anti-β-catenin ascites fluid (C7082, Sigma); rabbit Ab to α-catenin (ab2981, Abcam); H-90, rabbit Ab to p120 (sc-13957, Santa Cruz); mouse Ab to β-actin (8H10D10, Cell signalling Technology); mouse Ab to Phospho-Myosin Light Chain 2 (Ser19) (3671S, Cell Signaling Technology); mouse Ab to K14 (gift from Professor Leigh); Alexa Fluor 488 conjugated phalloidin for F-actin (A12379, ThermoFisher); Alexa Flour 488 conjugated rabbit Ab to non-muscle myosin IIa (ab204675, Abcam); D8H1X, rabbit Ab to YAP (D8H1X-XP, Cell Signaling Technology); EP1675Y, rabbit Ab to YAP1 (phosphor S127) (ab76252, Abcam); C-20, rabbit Ab to FOXM1 (sc-502, Santa Cruz); H-432, rabbit Ab to Cyclin A (sc-751, Santa Cruz); PC10, mouse Ab to PCNA (sc-56, Santa Cruz); Y69, rabbit Ab to c-Myc (ab32072, Abcam); 8H10D10, mouse Ab to beta actin (3700S, Cell Signaling Technology); mouse Ab to 14-3-3 gamma (Santa Cruz); rabbit Ab to LATS1+2 (ab70565, Abcam), normal rabbit IgG (2729S, Cell Signaling Technology); purified mouse IgG1(401401, Biolegend); 14C10, rabbit Ab to Glyceraldehyde-3-phosphate dehydrogenase (GAPDH)-Loading control (14c10, Cell Signaling Technology); B-6, mouse Ab to heat shock cognate 71 kDa protein (Hsc70)-Loading control (sc-7298, Santa Cruz); anti-Lamin A antibody (ab26300, Abcam); Secondary Abs were anti-mouse/rabbit IgG peroxidase antibody produced in goat (A0168/A6667; Sigma); Alexa Fluor 488 goat anti-mouse/rabbit IgG (A11029/A11034; Invitrogen) and Alexa Fluor 568 goat anti-mouse/rabbit IgG (A11031/A11036; ThermoFisher).

### 2.2. Cell culture and siRNA transfection

Various epithelial cell types were used in the study; HaCaT immortalized human skin keratinocyte line, SqCC/Y1 human oral buccal SCC line, a cutaneous squamous cell carcinoma T8 cell line (gift from Professor Harwood) with transduction of empty vector control (Vect Ct) and full length Dsg3 (FL) and primary keratinocytes derived from foreskin and breast skin and oral mucous tissues. HaCaT cell line was cultured in Dulbecco’s Modified Eagle Medium (DMEM; Lonza) supplemented with 10% fetal calf serum (FCS; 0200850, First Link (UK) Ltd) and SqCC/Y1 cells and primary keratinocytes were routinely maintained in EpiLife medium (with 60 mM calcium concentration) (MEPI500CA) supplemented with Human Keratinocyte Growth Supplement (HKGS; ThermoFisher Scientific). For the experiments, the media for SqCC/Y1 and primary cultures were replaced with modified Keratinocyte Growth Medium (KGM) (DMEM: Ham’s F12 = 3:1 supplemented by insulin (5 µg/ml), hydrocortisone (0.4 µg/ml), plus 10%FCS). T8-Vect Ct and D3 cell lines were cultured in KGM all time. A siRNA sequence specific for human Dsg3 mRNA, corresponding to nucleotides 620 to 640 of the respective coding region (Accession: NM_001944.1) (AAATGCCACAGATGCAGATGA) was designed and this sequence was subjected to a BLAST database search prior to being synthesized by Dharmacon Research (USA) [34,54]. All the other siRNA sequences were purchased from Dharmacon (USA) (ON-TARGETplus siRNA-2 J-011646-05, siRNA-3 J-011646-06, and siRNA-4 J-011646-08, Human DSG3, NM-001944), plus two scrambled controls, one was a randomized version of Dsg3 siRNA sequence (AACGATGATACATGACACGAG) and another, a randomized Dp siRNA (Scram-2: AACAGCGACTACACCAATAGA), all were synthesized and/or provided by the same company [34,54,58]. PKP1 siRNA (ON-TARGETplus PKP1 siRNA J-012545-05-0002 were purchased from Dharmacon (USA). Transient transfection with scrambled and Dsg3 specific siRNAs at the 100 nM concentration was conducted using the protocol as previously described [34,53,54]. For the generation of stable T8 Vect Ct and FL lines, the routine procedures were used as described previously [55].

### 2.3. Application of cyclic strain

The regimen for the cyclic strain was adapted from a previous publication [44]. Briefly, cells were plated and grown for 1∼2 days on collagen-coated BioFlex 6-well culture plates with flexible silicone elastomer bottoms (BF-3001C, Flexcell® International Corporation). Each plate was placed over the loading station containing 6 planar faced posts. Cell monolayers were subjected to equiaxial cyclic mechanical stretching with strain range of amplitude in 10∼15% and a frequency of 5 Hz, in a Flexcell FX-5000 Tension System (Flexcell International, Hillsborough, Burlington, NC) for varying durations. Control cells were seeded in the same BioFlex plates along with the strained cells but maintained at static state without any exposure to mechanical stretch. Lysates were extracted either immediately after a strain or transferred to static state and harvested later at different time points, for mRNA analysis by RT-qPCR or protein analyses by Western blotting and fluorescent microscopy.

### 2.4. Calcium-independent desmosome analysis

Cells seeded in BioFlex plates/Collagen I and subjected to cyclic strain or no strain. After that, cells were washed in calcium and magnesium-free Hanks’ balanced salt solution (14175053, HBSS, Gibco) briefly before being incubated in calcium-free medium (21068-028, calcium-free DMEM (Gibco) supplemented with 10% decalcified FCS plus 3 mM EGTA for 90 minutes, following the established protocol [28]. Control samples were cells incubated in growth medium with normal calcium concentration. Finally, all samples were fixed with 3.6% formaldehyde for 10 minutes only before processing for desmoplakin immunofluorescence.

### 2.5. Immunofluorescence, microscopy and image analysis

Cells either grown on BioFlex plates with or without cyclic strain or seeded on coverslips coated with polyacrylamide gel were washed briefly with PBS for a couple of times before fixation with either ice-cold methanol or 3.6% formaldehyde for 10 minutes at room temperature, according to the experiments. For total protein analysis, the formaldehyde-fixed samples were subjected to permeabilization with 0.1% Triton for 5 minutes before antibody staining. For peripheral protein analysis, cells were fixed with 3.6% formaldehyde without Triton permeabilization. The nonspecific binding sites in samples were blocked for 15-30 minutes with 10% goat serum in washing buffer before the primary and secondary antibody incubations, each for 1 hour at room temperature, respectively. For F-actin and Myosin IIa staining, Alexa Fluor 488 conjugated phalloidin or anti-Myosin IIa was incubated together with the secondary antibodies. All antibodies were diluted in 10% goat serum (G9023, Sigma). Coverslips were washed 3 times with washing buffer (PBS containing 0.2% Tween 20) after each antibody incubation. Finally, coverslips were counterstained with DAPI for 8–10 minutes before being mounted on microscope slides. Images of fluorescent staining were acquired with a Leica DM4000 epi-fluorescence microscope or confocal Zeiss710/880. All images were analyzed with ImageJ software using the default threshold algorithm for each channel across different samples/condition. Finally, total immunofluorescence intensity (IMF) per cell was calculated as the integrated density per channel divided by the total cell number (measured by the number of Nuclei in the DAPI channel). The data were normalized against the non-strained control in each experiment. For analysis of cytoplasmic and nuclear staining, the IMF intensity in each compartment was measured separately. This was achieved by subtracting the binary image of the nuclei from the channel of interest. The IMF was then measured on these new images with cytoplasmic signal only. The nuclear signal was calculated by subtracting total IMF by the cytoplasmic signal from each channel. The IMF intensity per cell for each compartment was calculated as described above. For the peripheral protein staining, the Nuclear mask was expanded using the ‘dilate’ tool. The remaining procedures were the same as described above.

### 2.6. Preparation and Functionalization of polyacrylamide (PA) gels

PA gels were prepared and modified from the previous work [20,38]. Briefly, coverslips were cleaned by sonication in ethanol and air-dried overnight. On the following day, all coverslips were incubated with a 5% 3-aminopropyltrimethoxysilane solution (281778, Sigma-Aldrich) in water for 30 minutes, followed by another 15 minutes incubation on a 0.5% Glutaraldehyde solution (A17876, Alfa Aesar) in PBS, both under agitation. Meanwhile, PA solutions were prepared with a ratio variation between 40% polyacrylamide (AAm) (1610140, Biorad) and 2% bis-acrylamide (Bis) (1610142, Biorad) which resulted in different modulus of the PA gels. In this study, we used three gel formulae; AAm 7.5%, 12%, 12% and Bis 0.05%, 0.145%, 0.45% respectively. Therefore, through micro-indentation tests, the results for the modulus of PA gels were approximately 8, 70 and 214 kPa respectively. Once prepared, the gel precursor solution was degassed for 5 minutes before gelation. To initiate gelation, 6 ul of 10% ammonium persulfate (APS; A3678, Sigma-Aldrich) and 4 ul of fN, N, N′, N′-Tetramethylethylenediamine accelerator (TEMED; T9281, Sigma-Aldrich) were added to 1 ml of the gel precursor solution. Six 10 ul drops of the final mixture were placed onto a Sigmacote® microscope slide. The activated coverslips were quickly placed on top of each drop, allowing them to flatten while adhering to the glass. Gels were left to polymerize at room temperature for 5 min. Afterward, gels and coverslips were flooded with a 50 mM HEPES solution (H3375, Sigma-Aldrich), gently pushed aside to the edge of the microscope slide and lifted from the surface. The coverslips were then stored in the same 50 mM HEPES solution until use. PA gels were functionalized through incubation with 0.2 mg/ml solution in water of the cross-linker Sulfo-SANPAH (2324-50, Biovision) and double 5 minutes irradiation by 365 nm light. This allowed the photoactivation of the cross-linker. Gels were washed with both deionized water and PBS. Lastly, gels were incubated with a 50 ug/ml collagen type I solution in PBS (354236, Corning) at room temperature for 3 hours. Prior to cell seeding, gels were sterilized with UVB irradiation and a 70% ethanol solution followed by a 30 minutes media incubation.

### 2.7. Western blotting, Co-immunoprecipitation, and Biotinylation assay

Total cell extraction, Triton-soluble and -insoluble fractionations, Western blotting and co-immunoprecipitation were performed following the protocols as described previously [54,55]. Protein concentrations in all samples were determined by Bio-Rad *DC* protein assay as a routine. For Western blotting, 5–10 µg of total proteins were resolved by SDS–PAGE and transferred to nitrocellulose membrane before antibody incubations. Equal loading was confirmed before probing for the target proteins in each set of samples. For co-immunoprecipitation (co-IP), ∼1 mg protein lysate per sample was used to immunoprecipitate YAP, pYAP and Dsg3 associated protein complexes, respectively, using 3–5 µg of primary antibody against each protein and protein-G magnetic Dynabeads (10003D, Invitrogen) and incubated overnight at 4°C on rotation. Finally, after washing thoroughly (4x in RIPA buffer and 1x in TTBS) the precipitate was re-suspended in 2xLaemmli sample buffer and heated for 3 minutes at 95°C before being resolved by SDS-PAGE followed by Western blotting procedures. For Dsg3 IP, calcium ion at the final concentration of 1.8 mM was added into the RIPA and washing buffers throughout the co-IP procedures. The extraction of surface proteins in strained and non-strained cells was performed by following the instruction provided by Pierce Cell Surface Protein Isolation Kit (Thermo Scientific).

### 2.8. Reverse transcription absolute quantitative RT-PCR (qPCR)

mRNA harvested using Dynabeads mRNA Direct kit (Invitrogen) was converted to cDNA using qPCRBIO cDNA Synthesis kit (#PB30.11-10, PCRBIO Systems, UK) and the cDNA was diluted 1:4 with RNase/DNase free water and stored at −20°C until used for qPCR. Relative gene expression qPCR was performed using qPCRBIO SyGreen Blue Mix Lo Rox (#PB20.11-50, PCRBIO Systems, UK) in the 384-well LightCycler 480 qPCR system (Roche) according to our well-established protocols [21] which are MIQE compliant [8]. Briefly, thermocycling begins with 95°C for the 30s prior to 45 cycles of amplification at 95°C for 1s, 60°C for 1s, 72°C for 6s, 76°C for 1s (data acquisition). A ‘touch-down’ annealing temperature intervention (66°C starting temperature with a step-wise reduction of 0.6°C/cycle; 8 cycles) was introduced prior to the amplification step to maximize primer specificity. Melting analysis (95°C for the 30s, 65°C for 30s, 65-99°C at a ramp rate of 0.11°C/s) was performed at the end of qPCR amplification to validate single product amplification in each well. Relative quantification of mRNA transcripts was calculated based on an objective method using the second derivative maximum algorithm (Roche). All target genes were normalized using a stable reference gene (POLR2A). The primer sequences for all genes analyzed in the study are listed in Supplementary Table 1.

### 2.9. Ki67 staining

The cells were fixed with 2% paraformaldehyde for 30 minutes at room temperature followed by permeabilization with 0.1% Triton for 5 minutes. Cells were stained with the anti-Ki-67 (M7240, Dako) and counterstained with DAPI. The proliferation rate was assessed by determining the Ki-67-expressing nuclei in relation to the total number of cells defined by DAPI staining [24].

### 2.10. Statistical Analysis

Statistical differences between controls and test groups were analyzed using unpaired, 2-tailed Student’s *t-test* in all cases. Data are presented as mean ± S.D. unless otherwise indicated. P values of less than 0.05 were considered statistically significant. Experiments were usually repeated three times. For image quantitation, all images were routinely acquired in 4-6 arbitrary fields per sample with a Leica DM4000 Epi-Fluorescence Microscope (upright) and analyzed with ImageJ software. For Western blotting analysis, lysates of three biologically independent replicates were collected from cyclic strain experiments. Wherever possible, the band densitometry of each blot was normalized against the loading control in the same lane, and the comparison between control and test group was normalized against the control and expressed as a fold change relative to control (set as 1).

## 3. Results

### 3.1. Mechanical cyclic strain alters junction protein expression and distribution in keratinocytes

To address our hypothesis that Dsg3 may respond to external mechanical loading in keratinocytes, we performed the equiaxial strain assay with skin-derived HaCaT and oral-mucosa derived SqCC/Y1 keratinocyte lines [51]. Cells were seeded at confluent densities in 6-well Flexcell plates for 1∼2 days prior to cyclic strain (15% amplitude, 5 Hz [44]) for approximately 6 or 24 hours. Control samples in this study were the cells seeded in the same plate without strain (stationary). First, immunofluorescence was performed to examine the expression and distribution of junctional proteins *in situ*, including Dsg3, using various fixation protocols which preferentially detect peripheral, total and cytoskeletal-associated proteins, respectively [55] (see Materials and Methods). We focused more on the peripheral protein analysis in many cases as it reflects the surface junction assembly. Since E-cadherin and nonmuscle actomyosin have been identified as force sensors [5,12,14,15,29,32], we monitored the expression of E-cadherin and Myosin IIa alongside Dsg3 in both cell lines (Fig. 1*A*, Fig. S1). It was found that the peripheral protein analysis (-Triton) was more revealing than the other two protocols and showed a marked increase of both cadherins and Myosin IIa in their response to mechanical cyclic strain, in particular, in SqCC/Y1 cells, compared to their static counterparts (Fig. 1*A*, or data not shown). The peripheral protein expression of Dp and α-catenin also showed an evident increase in strained SqCC/Y1 cells (Fig. S1). Compared to SqCC/Y1, we noticed fewer alterations in HaCaT cells except for pMLC which exhibited a clear rise in strained cells compared to stationary control (Fig. S1 & S2). Next, Triton-soluble (cytoskeletal non-associated) and insoluble (cytoskeletal-associated) fractions, were collected separately and analyzed by Western blotting as described previously [54,55]. We detected strain induced elevation of most junctional proteins, including E-cadherin and Dsg3, in both fractions in SqCC/Y1 cells and again, to a lesser extent, in HaCaTs (Fig. 1*B*, Fig. S3). Together, these results demonstrated that environmental mechanical loading induces increased expression and surface assembly of junctional proteins, including Dsg3 and E-cadherin, with changes more evident in oral keratinocytes than skin-derived HaCaT cells.

**Fig. 1.**
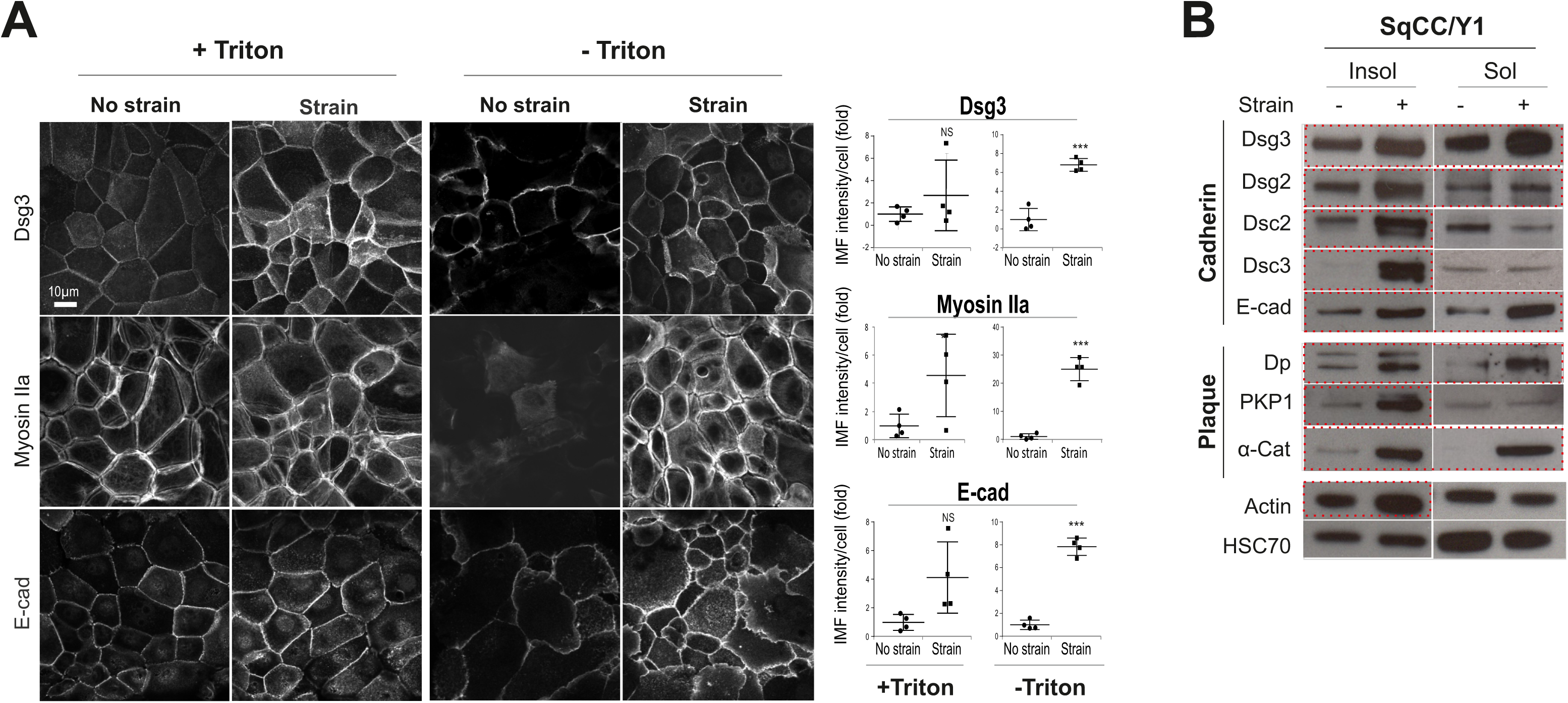
Cyclic strain causes enhanced Dsg3 and E-cadherin expression and their surface assembly in oral keratinocytes. *A*, Confocal microscopy of SqCC/Y1 cell line plated in Flexcell wells at confluent density for 1 day before being subjected to cyclic strain or no strain, for 6 hours prior to fixation with formaldehyde and then permeabilization with Triton X-100 (+Triton) or no permeabilization (-Triton). Samples were immunostained for the indicated proteins. Cells fixed with formaldehyde only detected preferentially the peripheral protein expression and this differed from cells treated with Triton that removed some soluble proteins. Images were the projections of confocal stacks with maximal intensity. Quantitation for each protein is shown on the right (n>4, mean ± S.D., NS: no significance, *p<0.05, ***p<0.001). Scale bar is 10 µm. *B*, Western blotting analysis for junctional proteins in subcellular fractions, *i.e.* Triton-soluble (Sol) and insoluble pools (Insol), in SqCC/Y1 cells subjected to no strain (-) and strain (+) for 6 hours. HSC70 was used as a loading control. Those marked by the red dotted boxes indicate an evident increase in the expression in strained relative to static cells in either fraction, respectively.

### 3.2. Dsg3 Is Required for E-cadherin and Actomyosin Junction Assembly in Response to Mechanical Strain

To address the importance of Dsg3 in response to cyclic strain, we performed a Dsg3-knockdown study with transient transfection of siRNA (100nM) in HaCaTs in conjunction with cyclic strain. Both RT-qPCR and Western blotting analyses were performed in cells harvested instantly after strain or no strain, validating significant knockdown of Dsg3 at the transcript and protein levels (Fig. S4). No reduction for other desmosomal cadherins, including *DSG2*, was found at the transcript levels as well as their response to the strain, except for *DSP* that showed an increase in strained cells with Dsg3 depletion compared to its static counterpart (Fig. S4*A*). On the other hand, *DSC2*, as well as Dsc2 protein, showed a significant increase following Dsg3 depletion compared to controls, suggesting its likely compensation for the loss of desmosomal cadherins, such as Dsc3 and Dsg2 in addition to Dsg3 (Fig. S4*B*). Notably, PKP1 displayed an evident decrease in the Triton insoluble fraction in knockdown cells. In addition, an increase of actin in the insoluble fraction with a diminution of its soluble counterpart was also observed in cells in response to strain (Fig. S4*B*). It is worth noting that a minor increase of residual Dsg3 frequently was observed in the knockdown cells subjected to strain suggesting its sensitivity to mechanical loading as described above. Nevertheless, some variations were observed in different protein fractionation analysis in this study. Significantly, immunostaining for peripheral E-cadherin and Myosin IIa revealed their marked reduction, with disruption of E-cadherin at the junctions, in Dsg3 knockdown cells with exposure to cyclic strain (arrow Fig. 2*A,B*), indicating that Dsg3 depletion had a negative impact on E-cadherin junction assembly in its response to mechanical loading. In support, analysis of T8 cell lines with Dsg3 overexpression (T8-D3) (gain-of-function) demonstrated enhanced E-cadherin and Myosin IIa at the junctions as opposed to T8-Vector control line (Vect Ct) or T8-D3 cells with Dsg3 RNAi that evoked disruption of junctions (Fig. S5). Together, these results demonstrated that Dsg3 is required for junction assembly in keratinocyte response to mechanical loading.

**Fig. 2.**
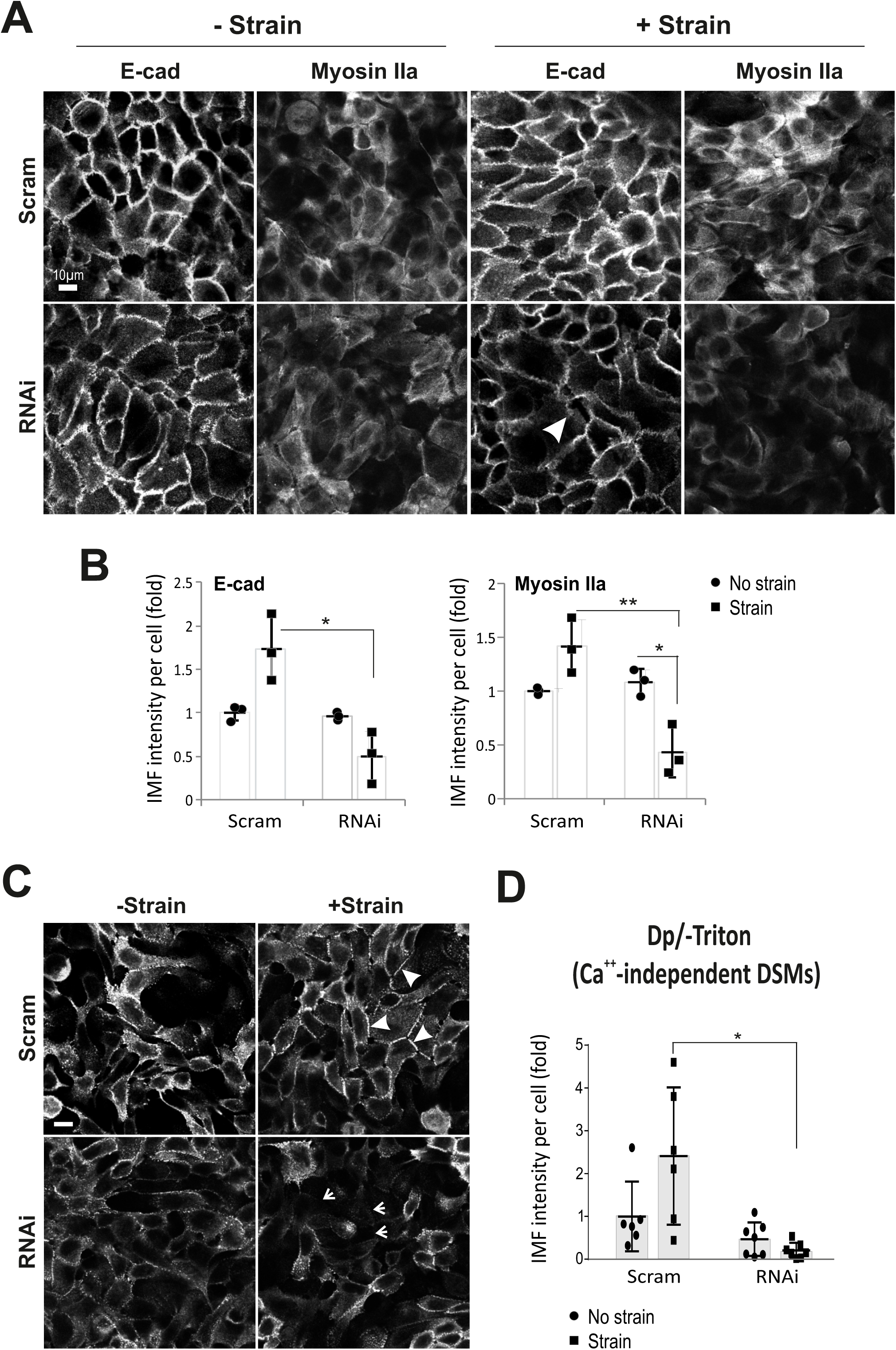
Dsg3 knockdown causes a defect of junction assembly in E-cadherin and Myosin IIa as well as the calcium-independent DSMs. *A*, Confocal images of E-cadherin (E-cad) and Myosin IIa staining in HaCaT cells subjected to strain (+Strain) and no strain (-Strain), without (Scram) and with Dsg3 knockdown (RNAi), respectively. Cells were strained for 4 hours before being transferred to stationary, without changing the medium, for 90 minutes prior to fixation and immunostaining. *B*, Image quantitation for E-cadherin and Myosin IIa staining acquired with an Epi-fluorescent microscope (n=3, mean ± S.D., *p<0.05, **p<0.01). *C*, Confocal images of HaCaT cells subjected to strain, or no strain, for 4 hours. Cells were transferred to stationary after strain and incubated with a calcium-free medium for 90 minutes in an incubator prior to fixation (formaldehyde) and immunostaining for Dp, an established protocol for the analysis of calcium-independent DSMs [28]. Arrowheads indicate the linear Dp staining at the junction and arrows show cells missing of the calcium-independent DSMs. *D*, Image quantitation for Dp staining (n=5∼7, mean ± S.D., pooled from two experiments, *p<0.05). Scale bar is 10 µm.

DSMs confer strong cell-cell adhesion by forming calcium-independent junctions in confluent keratinocyte cultures [28]. In this regard, we asked whether mechanical strain promotes the formation of calcium-independent DSMs and thus we analyzed the nature of DSMs by adopting the established protocol for Dp staining with minor modification [28]. After cyclic strain, cells were incubated in a calcium-free medium for 90 minutes, prior to fixation and immunostaining for Dp. The result showed numerous calcium-independent DSMs in HaCaT cells of all conditions. In control siRNA treated cells, the mechanical strain resulted in a more even and concentrated linear distribution of Dp at the junctions (arrowheads Fig. 2*C*). In contrast, Dsg3 knockdown resulted in punctate Dp staining pattern in non-strained cells but with a marked reduction in strained cells compared to the respective controls (arrows Fig. 2*C,D*). This finding indicates a defect in the DSM assembly induced by cyclic strain in Dsg3 depleted cells and reinforces that Dsg3 is required for both AJ and DSM junction assembly in response to mechanical force. Furthermore, the protein turnover for various junctional proteins, including Dp, in response to strain for up to 24 hours was analyzed and the result showed a gradual decline of Dp in response to strain compared to non-strained cells (Fig. S6), indicating dynamic remodeling of the DSMs in response to mechanical loading. Subtle changes were observed in Dsg3, E-cadherin and Pg (γ-catenin) implying potentially a large intracellular store of redundant proteins whose presence masks detection of protein turnover in this experiment. Collectively, these results suggest that mechanical strain does not necessarily enhance the formation of calcium-independent DSMs but rather causes alterations in protein stability, *e.g.* Dp, and junction remodeling.

### 3.3. Dsg3 Regulates the Expression and Localization of YAP in Response to Mechanical Loading

YAP has been identified as a mechanosensor [39] with nuclear relocation of YAP being regulated by mechanical strain [24] and substrate stiffness [41]. To elucidate the underlying mechanism of how Dsg3 regulates junction formation induced by mechanical loading, we investigated YAP and pYAP-S127 (pYAP) expression in a time-course experiment in HaCaTs with exposure of strain for 6 hours. In this case, cells were transferred to the stationary state after straining (relaxation) in an incubator and total lysates were then extracted at various time points at 0, 2, 6, 24, 48, and 72 hours, alongside with static control cells (Fig. 3*A*). Consistently, no evident increase was observed in Dsg3 in samples harvested immediately after strain compared to a stationary sample (0h, Fig. 3*B*). However, a gradual increase in Dsg3 was detected later in post-strained cells, compared to non-strained, from 6 hours with a peak at 48 hours. Interestingly, in response to cyclic strain, pYAP showed an initial reduction compared to static cells (0h) and this was followed by a swift recovery within 2 hours in post-strained cells, with climbing up to the baseline of static control cells and maintained at this level for up to 72 hours of the experiment (Fig. 3*B,C*). YAP also showed a trend with an increase in post-strained cells. In contrast, in static populations, both pYAP/YAP showed a decline after 24 hours and remained at this low level until 24/48 hours, indicating their degradation in stationary confluent cultures. Generally, it was observed that these three proteins exhibited a similar shift in their response to mechanical loading, with a gradual increase and/or stabilization at the baseline compared to static cells in which the activity of pYAP (indicative Hippo) appeared to be transient and was gradually switching off in mature and well-established confluent cultures. To confirm such a delayed response for pYAP/YAP, the strained cells were transferred to stationary in an incubator, and either fixed or extracted 24 hours later after strain. Immunostaining for peripheral pYAP indicated a striking increase along with Dsg3 in post-strained cells in contrast to static counterpart (Fig. 3*D*) and Western blotting also confirmed increased expression of pYAP/YAP in post-strained cells compared to static control (data not shown).

**Fig. 3.**
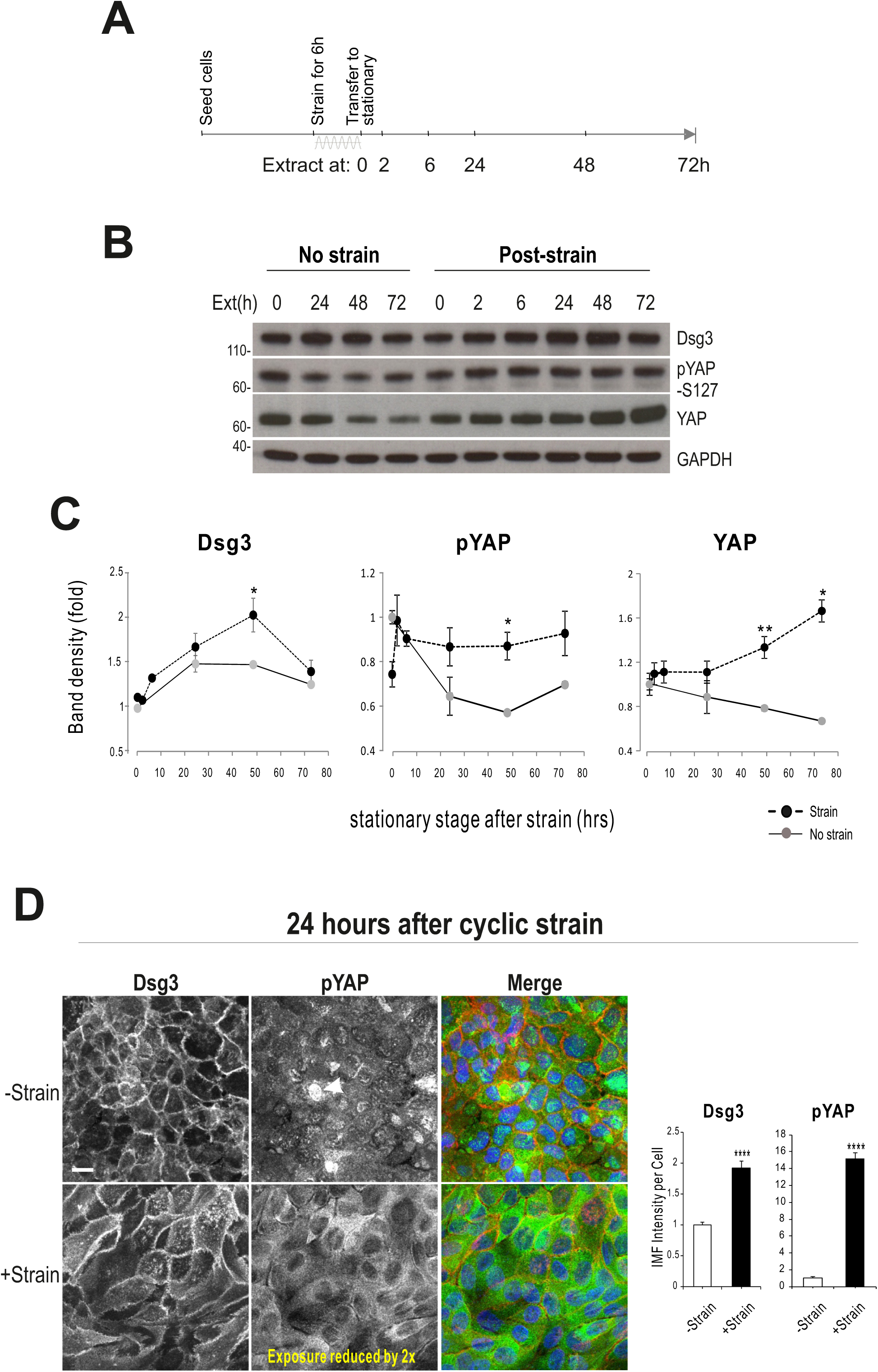
The expression of YAP and pYAP appeared to coincide with Dsg3 in response to cyclic strain. *A*, Timeline of a cyclic strain experiment. HaCaT cells were strained, or non-strained, for 6 hours before being extracted either immediately (0h) or later, after being transferred to stationary, at various time points as indicated. *B&C*, Western blotting and band densitometry analysis of Dsg3, pYAP and YAP expression. GAPDH was used as a loading control. Note that all three proteins showed increased expression in response to cyclic strain compared to non-strained samples. Dsg3 exhibited an increase with retardation and reached the maximum 48 hours later after strain. pYAP showed an initial reduction after strain (0h) that was followed by swift recovery to the baseline of non-strained cells (0h) and maintained at this level until the end of this experiment (72h). In contrast, pYAP in non-strained cells was falling over time. YAP, however, showed increasing in strained and decreasing in static control cells. Quantitation for each protein was normalized against the loading in each lane and presented as fold change of the non-strained sample at 0h time point (n=3, mean±S.E.M, *p<0.05, **p<0.01). *D*, Immunofluorescence (-Triton) in post-strained and static cells 24 hours after strain. Note that the exposure of the confocal image in +strained sample was reduced two times. (n=5, mean±S.D. ****p<0.0001). Scale bar, 10 µm.

Next, immunofluorescence for YAP and pYAP was performed that revealed both proteins were highly soluble to non-ionic detergents, such as Triton. Therefore, we focused on the peripheral protein analysis (-Triton) in HaCaTs that were subjected to Dsg3 knockdown in conjunction with cyclic strain using the same regimen. YAP staining in control siRNA treated cells displayed predominant cytoplasmic distribution and the mechanical strain resulted in its elevation (∼2-fold, Fig. S7*A*). In contrast, Dsg3 depletion resulted in a marked decrease of YAP with concomitantly enhanced nuclear translocation in response to strain (Fig. S7, Fig. 4*C*). Intriguingly, the pYAP staining revealed a sub-pool of protein located at the cell borders, with a trend of increase in cells exposed to mechanical loading (control cells, Fig. 4*A*). In contrast, the cyclic strain of Dsg3 depleted cells evoked a remarkable reduction of pYAP, with concomitant nuclear translocation, compared to static counterparts (Fig. 4*B,C*), that was confirmed by Western blotting analysis (Fig. 5*A*). Immunofluorescence analysis for pYAP in various non-keratinocyte cell types revealed that the membrane pYAP was specific in keratinocytes since the other cell types, such as oral fibroblasts, 3T3, and Cos-1 cells, showed no evident pYAP membrane staining except for NTERT cells which also exhibited higher pYAP levels than other cell types (Fig. S8).

**Fig. 4.**
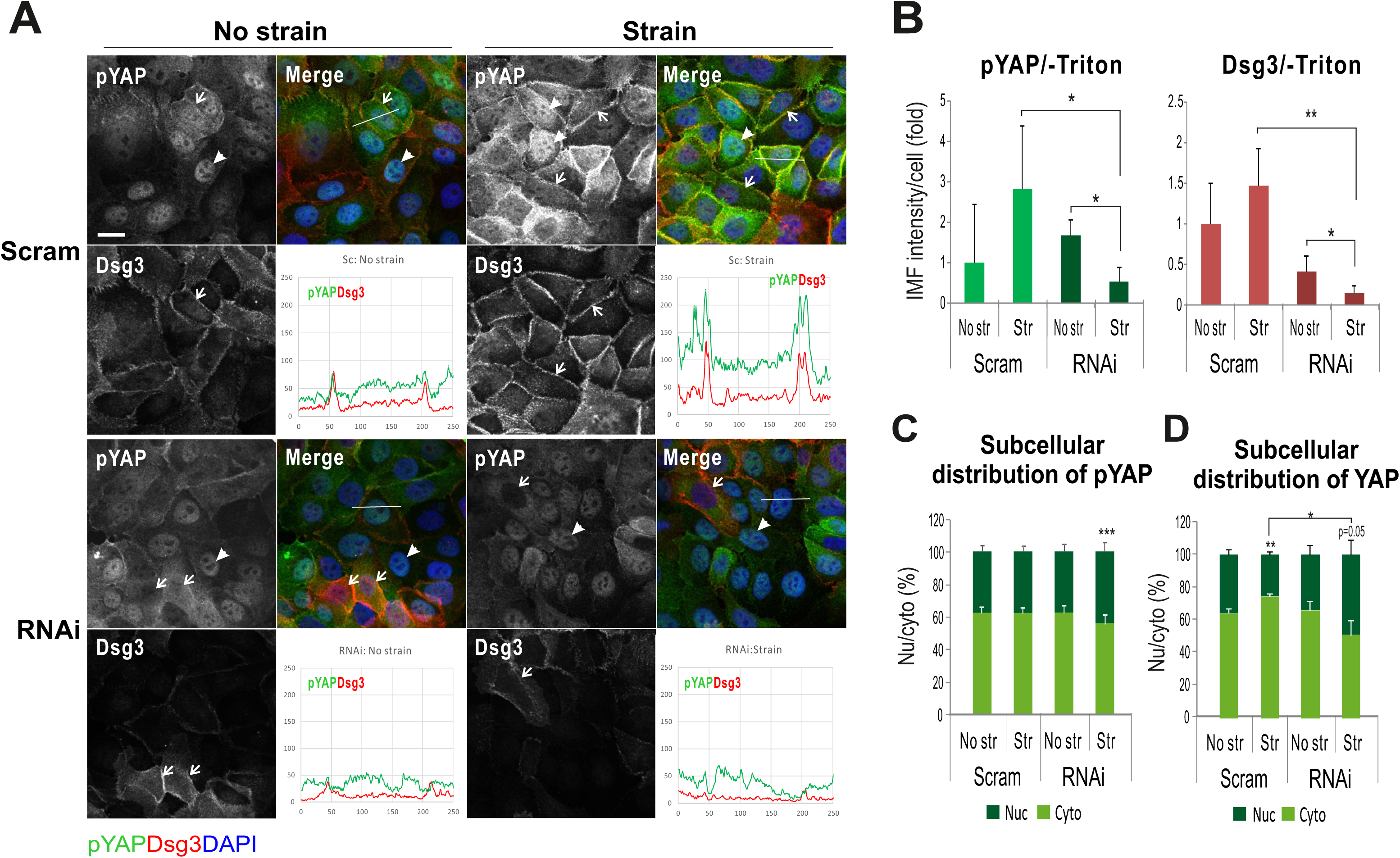
Dsg3 regulates peripheral pYAP expression and localization in its response to cyclic strain. *A*, Confocal images of HaCaT cells doubled labeled for Dsg3 (red) and pYAP (green). The siRNA pre-treated cells (Scram: control siRNA; RNAi: Dsg3 specific siRNA) were seeded at confluent density in BioFlex wells for 1 day before subjected to cyclic strain for 6 hours or maintained in stationary condition prior to fixation with formaldehyde only and then immunostained for the indicated proteins. Arrowheads indicate pYAP nuclear staining and arrows indicate the cells which had Dsg3 expression with concomitantly less nuclear pYAP signals. Note the enhanced colocalization of pYAP and Dsg3 was shown in control cells subjected to strain and this was largely abrogated in cells with Dsg3 knockdown, in particular with strain (see the line profile in each condition). *B*, Image quantification for pYAP and Dsg3 expression and both channels were subject to nuclear extraction before measurement with a high threshold that detected mainly the peripheral signals. *C,D*, Subcellular distribution analysis for pYAP and YAP, respectively. (n=4, mean ± S.D., *p<0.05, **p<0.01, ***p<0.001). Scale bar, 10 µm.

**Fig. 5.**
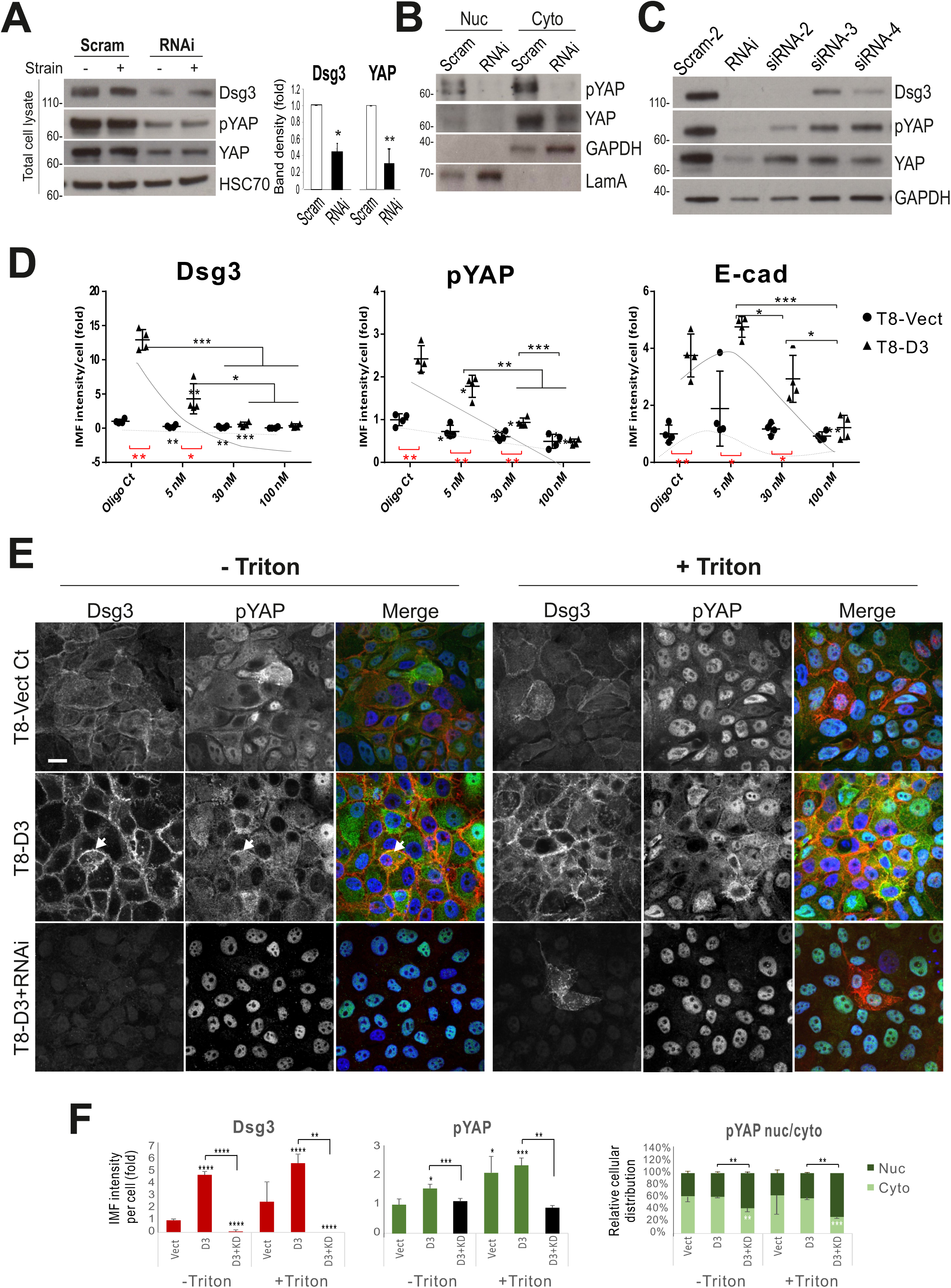
Effect of various siRNA-mediated knockdown or overexpression of Dsg3 on YAP and pYAP expression. *A*, Western blotting analysis for the indicated proteins in total lysates of HaCaT cells transfected with siRNA for 2 days. The bar charts show the band densitometry for Dsg3 and YAP blots (n=3, mean ± S.D.). *B*, Nuclear versus cytoplasmic fractionation analysis for pYAP and YAP. GAPDH and LamA were used as loading controls. *C*, Western blots of HaCaT cells transfected with various Dsg3 siRNA sequences (siRNAi: used for the majority of the experiments) alongside a second scrambled control siRNA (Scram-2). *D*, Titration of Dsg3 siRNA in cutaneous keratinocyte T8-Vect Ct and D3 (Dsg3 overexpression) cell lines by immunofluorescence. The siRNA pretreated cells were seeded on coverslips for 1 day before fixation and immunostaining for the indicated proteins. Only the image quantitation data were shown (n=4 fields/>350 cells were analyzed per sample). *E*, Confocal images of peripheral (-Triton) and total (+Triton) protein staining for Dsg3 (red) and pYAP (green) in T8 cell lines alongside T8-D3 cells with Dsg3 knockdown (T8-D3+RNAi). Arrows indicate colocalization of Dsg3 and pYAP in T8-D3 cells. Dsg3 knockdown in T8-D3 cells caused pYAP nuclear localization. Scale bar is 10 µm. *F*, Image quantitation of Dsg3 and pYAP expression as well as the subcellular distribution for pYAP (n=6, mean ± S.D.). (*p<0.05, **p<0.01, ***p<0.001, ****p<0.0001).

To validate the above finding, Dsg3 knockdown was performed with additional different siRNAs (ON-TARGETplus siRNA, Dharmacon) alongside a different scrambled siRNA control and again, a similar finding with a reduction of YAP/pYAP was observed in all knockdown cells compared to controls (Fig. 5*B*). Immunostaining in two different siRNA pre-treated cells seeded on coverslips at high and low densities also showed the nuclear retention of pYAP/YAP, with augmented nuclear/cytoplasm ratios, in sparse cultures in Dsg3 depleted cells in contrast to their confluent counterpart or control cells plated at low density (Fig. S9). Moreover, a titration of Dsg3-siRNA at a series of concentrations, *i.e*. 5, 30 and 100 nM, was conducted in our generated stable keratinocyte lines, T8-Vect control, and T8-D3. A dose-dependent response in pYAP reduction was detected in cells treated with various siRNA concentrations, but with compensation by elevated Dsg3 levels in T8-D3 cells (Fig. 5*C*). In line with enhanced peripheral pYAP in response to strain (Fig. 4), increased pYAP was also detected at the cell borders in T8-D3 compared to T8-Vect control and Dsg3 depletion abolished pYAP signals at the cell borders with a result of its marked nuclear relocation (Fig. 5*D,E*). In addition, E-cadherin also was analyzed here and showed increased expression in T8-D3, relative to T8-Vect control, but in sharp contrast, with attenuation in cells with Dsg3 knockdown, in a siRNA dose-dependent manner (Fig. 5*C*).

Taken together, these results support our hypothesis that Dsg3 regulates YAP/pYAP and is required for their nuclear export and the surface recruitment of pYAP in keratinocytes that might facilitate the AJ junction formation and the onset of CIP.

### 3.4. Dsg3 Colocalizes and Forms a Complex with pYAP which is Sequestered to the Plasma Membrane

Confocal microscopy further demonstrated the colocalization of Dsg3 and pYAP at the plasma membrane in HaCaT, as well as in primary, keratinocytes (Fig. 6*A*). Since YAP and pYAP were found to be dynamic proteins we monitored their expression profiles alongside Dsg3 in HaCaTs at sub-confluence, newly established-confluence, and over-confluence culture (up to 5 days) by Western blotting analysis. The expression of pYAP/YAP appeared to be transient and reached a maximum in freshly confluent cultures, similar to Dsg3, before decline when cells reached over-confluence (Fig. 6*B*). E-cadherin, Dp, PKP1/3, and Pg were also analyzed. While E-cadherin and PKP1/3 showed a steady increase, with E-cadherin peaked at fresh confluence and staying at the same levels for up to 5 days of the experiment, Dp reached a peak in fresh confluence but then underwent degradation with multiple degraded bands existing in over-confluent culture. Pg, however, showed less change in all aged cultures. Together, these data suggest that the regulation of Dsg3 and pYAP seems to be associated with one another but is independent of other junctional proteins analyzed, including E-cadherin. Next, co-immunoprecipitation (IP) assay with the antibody to pYAP-S127 as well as YAP, was performed in HaCaT lysates extracted from freshly confluent cultures. It was demonstrated that Dsg3 co-immunoprecipitated with pYAP indicating they form a complex, but yet was barely detectable for YAP (Fig. 6*C*). IP with anti-Dsg3 initially failed to detect pYAP but later, a diffuse band of pYAP was detected in conditions with calcium addition in buffers throughout the IP procedures. To confirm such a complex formation in primary keratinocytes, co-IP was repeated in foreskin cells and in this case, Dsg3, as well as PKP1/3 and 14-3-3, were detected in the complex purified with anti-pYAP IgG and only PKP1 and 14-3-3 were found in the YAP IP (Fig. 6*D*). These findings demonstrated that Dsg3 forms a complex with pYAP and helps to recruit it to the cell surface via a protein complex containing PKP. Since NTERT cells exhibit the most striking pYAP membrane distribution (Fig. S8), we knocked down Dsg3 in this cell line and examined the impact on pYAP peripheral distribution. Double immunostaining for pYAP/Dsg3 demonstrated the loss of pYAP at the plasma membrane with reduced peripheral cytoplasmic expression in Dsg3 depleted cells compared to control (Fig. 7*A,B*). To determine the specific role of PKP1 in this process, we performed PKP1 knockdown in HaCaT cells and double labeled pYAP/Dsg3 for fluorescent microscopic examination (Fig. 7*C*). It was revealed that PKP1 silencing caused disruption of Dsg3 at the junctions with concomitant loss of peripheral pYAP as well as reduced cytoplasmic staining, indicating a dependence of the Dsg3/pYAP interaction on PKP1 (Fig. 7*C,D*). Diffuse cytoplasmic Dsg3 staining with enhanced cell spreading like the Dsg3 knockdown cells [35] was also observed in cells with PKP1 depletion (Fig. 7*C*). Finally, biotinylated assay was performed to analyze the surface protein expression (Pierce kit, Thermo Scientific) in strained and non-strained cells and the results indicated enhanced expression for both cadherins as well as YAP/pYAP in strained versus non-strained cells (Fig. 6*E*). Taken together, these data suggest that Dsg3 connects to pYAP via PKP (see model below).

**Fig. 6.**
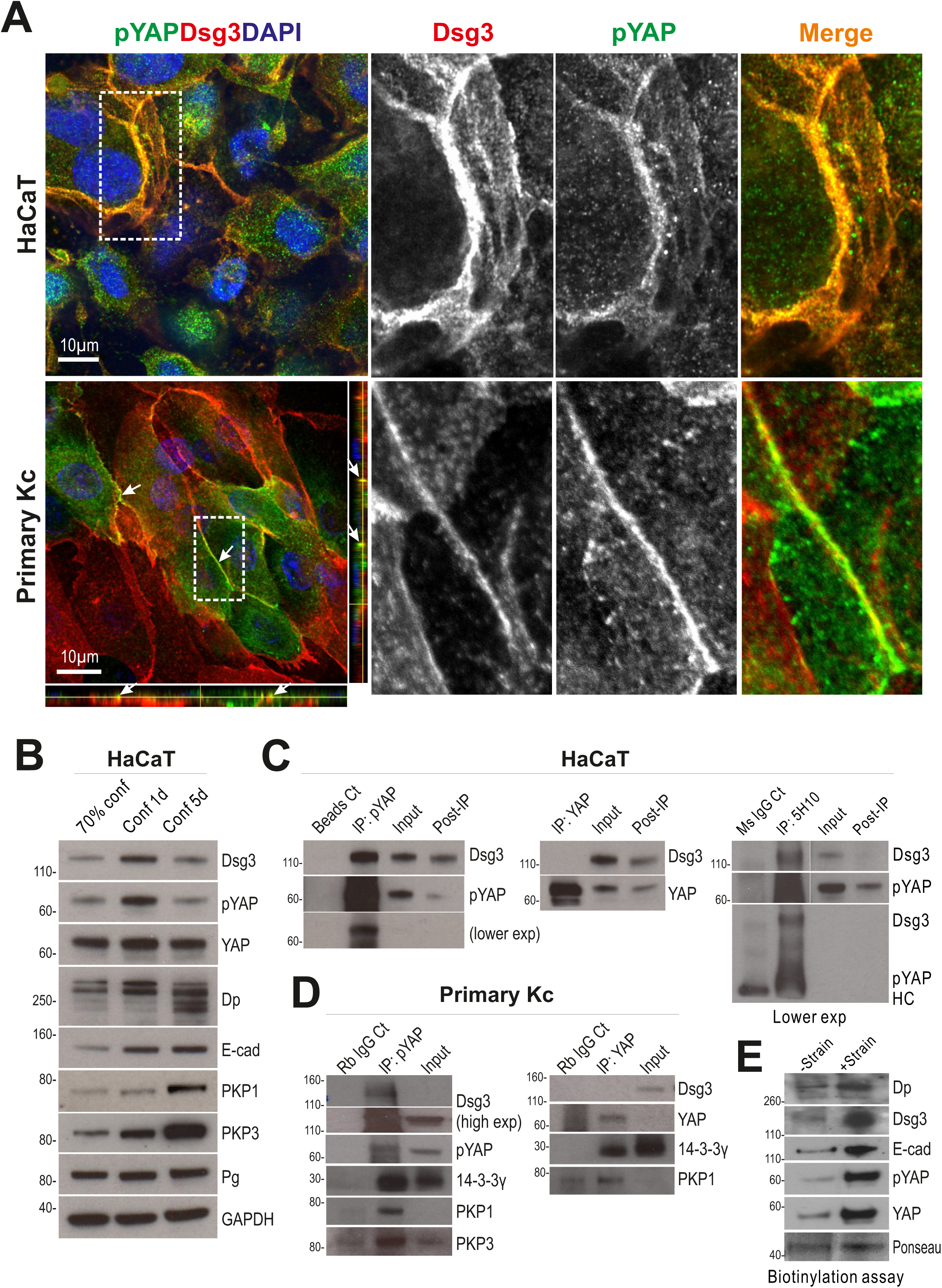
Dsg3 colocalizes and forms a complex with pYAP. *A*, Confocal super-resolution images of HaCaT and primary foreskin keratinocytes double labeled for Dsg3 and pYAP. The enlarged images of the white dotted boxes with merged and each channel are displayed on the right for each cell line. Arrows indicate the colocalization of both proteins. Scale bar is 10 µm. *B*, Western blotting of the indicated proteins in sub-confluent (70%), freshly-confluent (100% for 1d) and over-confluent cultures (5d) of HaCaT cells. GAPDH was used as a loading control. *C*, Co-IP analysis of immunoprecipitates in freshly confluent HaCaT cells, purified with anti-pYAP, anti-YAP, and anti-Dsg3 antibodies, respectively. The control lanes included Bead only, mouse IgG alone, input and post-IP samples. Lower exp: lower exposure. The input and post IP lanes on the right were with high exposure. *D*, Co-IP analysis of immunoprecipitates of primary foreskin keratinocytes, purified with anti-pYAP and anti-YAP antibodies, respectively. The control lanes were rabbit IgG alone and input before IP. *E*, Biotinylated assay for surface protein expression in non-strained and strained cells. HaCaT cells were seeded in Flexcell wells for 1 day before being subjected to strain or no strain for 5 hours. Then, the surface proteins were extracted using the Pierce Cell Surface Protein Isolation Kit. Ponceau staining was used as a loading control here.

**Fig. 7.**
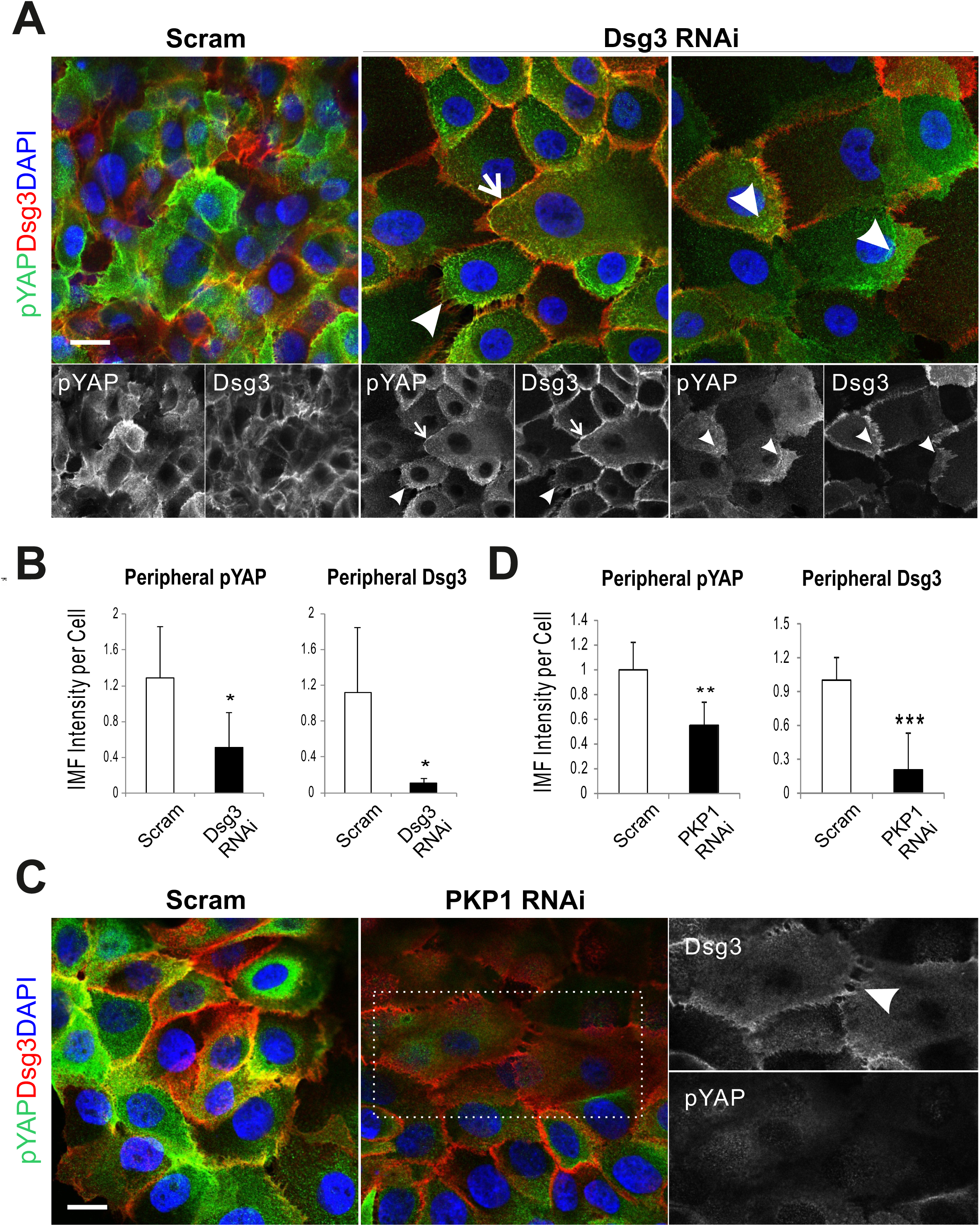
Either Dsg3 or PKP1 knockdown results in a reduction of peripheral pYAP expression. *A*, Confocal images of pYAP (green) and Dsg3 (red) staining in NTERT cells (-Triton) transfected with control or Dsg3 specific siRNA for 2 days. Arrows indicate the colocalization of both proteins at the plasma membrane. Arrowheads indicate the disruption of Dsg3 coupled with the loss of peripheral pYAP staining in the sub-membrane areas. *B*, Image quantitation of pYAP shown in A. *C*, Confocal images of pYAP (green) and Dsg3 (red) staining in cells transfected with control or PKP1 siRNA for 2 days. The enlarged images of the white dotted box for each channel are displayed on the right. Arrow indicates Dsg3 disruption at the junctions with concurrent loss of peripheral pYAP. *D*, Quantitation for cytoplasmic pYAP and Dsg3 expression. (n=4∼6 fields, mean ± S.D., *p<0.05, **p<0.01, ***p<0.001). Scale bar is 10 µm.

### 3.5. Elevated Expression of Dsg3 in Response Substrate Stiffness, similar to other Force Sensors

In addition to mechanical loading, substrate stiffness also is known to have a substantial impact on a wide range of cell behaviors, such as adhesion, spreading, locomotion, differentiation and cell fate decision [47,61]. It has been shown that E-cadherin serves as a mechanosensor in response to substrate stiffness [5,12,26]. To address whether Dsg3 also is responding to substrate stiffness, we examined Dsg3 expression, alongside E-cadherin and other well-known mechanosensors such as Myosin IIa and α-catenin, in cells seeded on collagen type I coated polyacrylamide hydrogels of varying stiffness (elastic modulus at approximately 8, 70, 215 kPa) that cover the biophysical stiffness of the skin [25], alongside glass coverslips (65 GPa [61]). Immunostaining for both cadherins, α-catenin, Myosin IIa, and F-actin indicated their expression correlated with substrate stiffness (Fig. 8*A*, Fig. S10). As anticipated, staining for YAP revealed enhanced nuclear localization in Dsg3 knockdown cells in a stiffness-dependent manner (Fig. 8*B,C*).

**Fig. 8.**
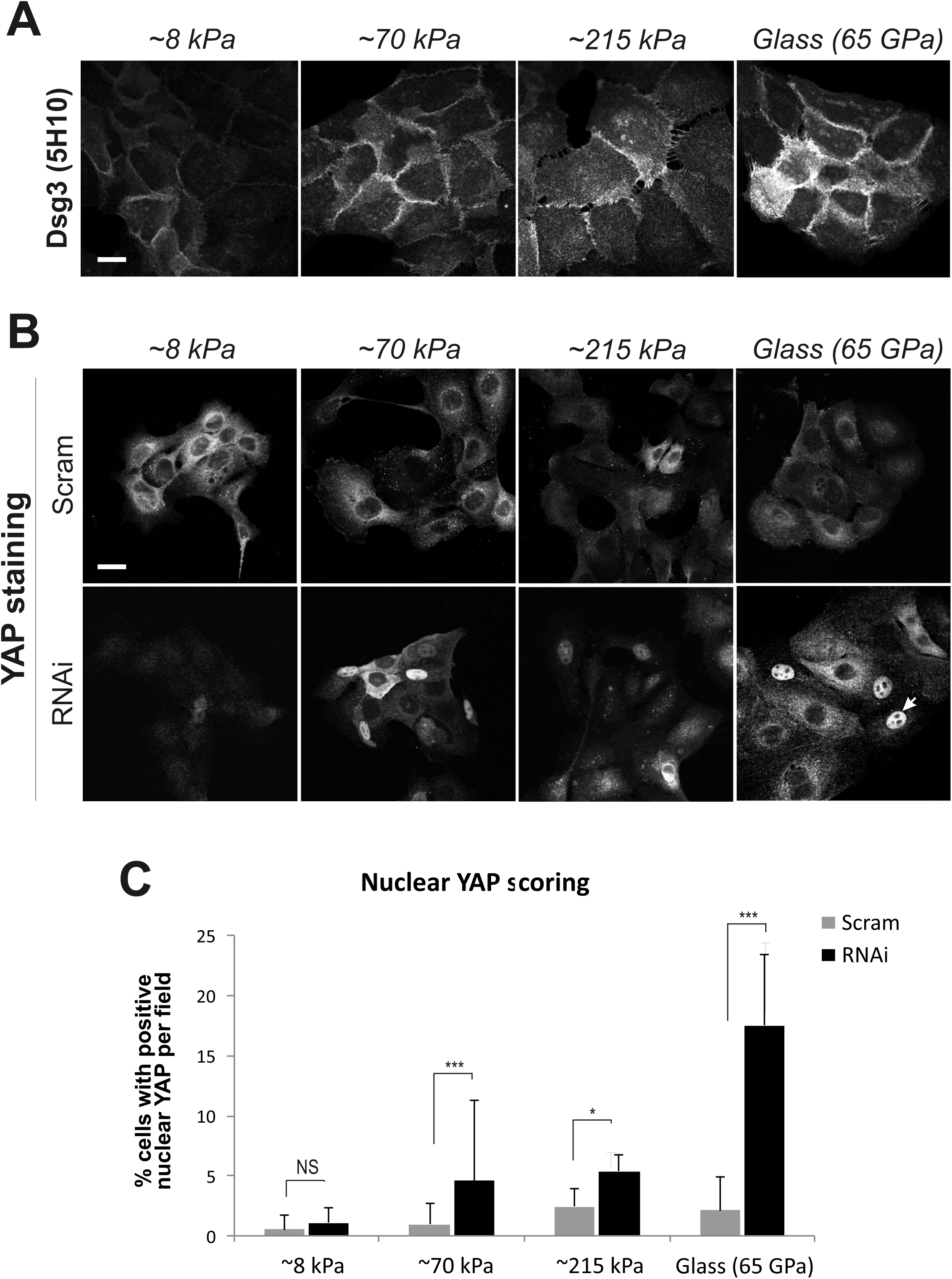
Dsg3 knockdown causes retardation of YAP nuclear exclusion in a substrate stiffness-dependent manner. *A*, Confocal images of Dsg3 staining in HaCaT cells seeded on collagen type I coated polyacrylamide hydrogels of varying stiffness along with glass coverslip. A correlation between protein expression and substrate stiffness was shown. *B*, Confocal images of YAP staining in siRNA pre-treated cells seeded on polyacrylamide hydrogels. *C*, Quantitation for the percentage of cells with positive YAP nuclear staining (n>8, *p<0.05, ***p<0.001, NS: no significance). Scale bar is 10 µm.

### 3.6. Dsg3 Knockdown Has an Impact on YAP Target Genes and cell proliferation

Transcript expression of several YAP target genes, such as *CYR61, CNGF, FOXM1, CMYC, CCNA2*, and *CENPA* [18,22,62] in HaCaT cells with/without Dsg3 knockdown and the results showed a marked reduction in the majority of these genes, including *YAP1* in Dsg3 depleted cells compared to control (Fig. 9*A*). Ki67 staining also indicated a significant reduction in Dsg3 knockdown cells (Fig. 9*B*). However, cyclic strain did not show any evident induction of these genes, in our current setting (confluent cells harvested immediately after strain), by qPCR analysis (data not shown) as well as in Ki67 staining in strained and post-strained cells for up to 3 days compared to the respective controls (Fig. S11). Nevertheless, Western blotting in lysates of strained and non-strained cells revealed an increase in FOXM1 (1.6-fold) and c-Myc (1.9-fold) in post-strained cells at 24 hours as compared to the respective static controls (Fig. 9*C*). Importantly, Dsg3 knockdown attenuated such responses in post-strained cells (2-fold reduction in FOXM1 and Cyclin A at 24 hours, Fig. 9*D*). Collectively, these results suggest that Dsg3 plays a role in regulating *YAP1* and its targeted gene expression although the cyclic strain had little effect on cell proliferation in confluent keratinocyte cultures.

**Fig. 9.**
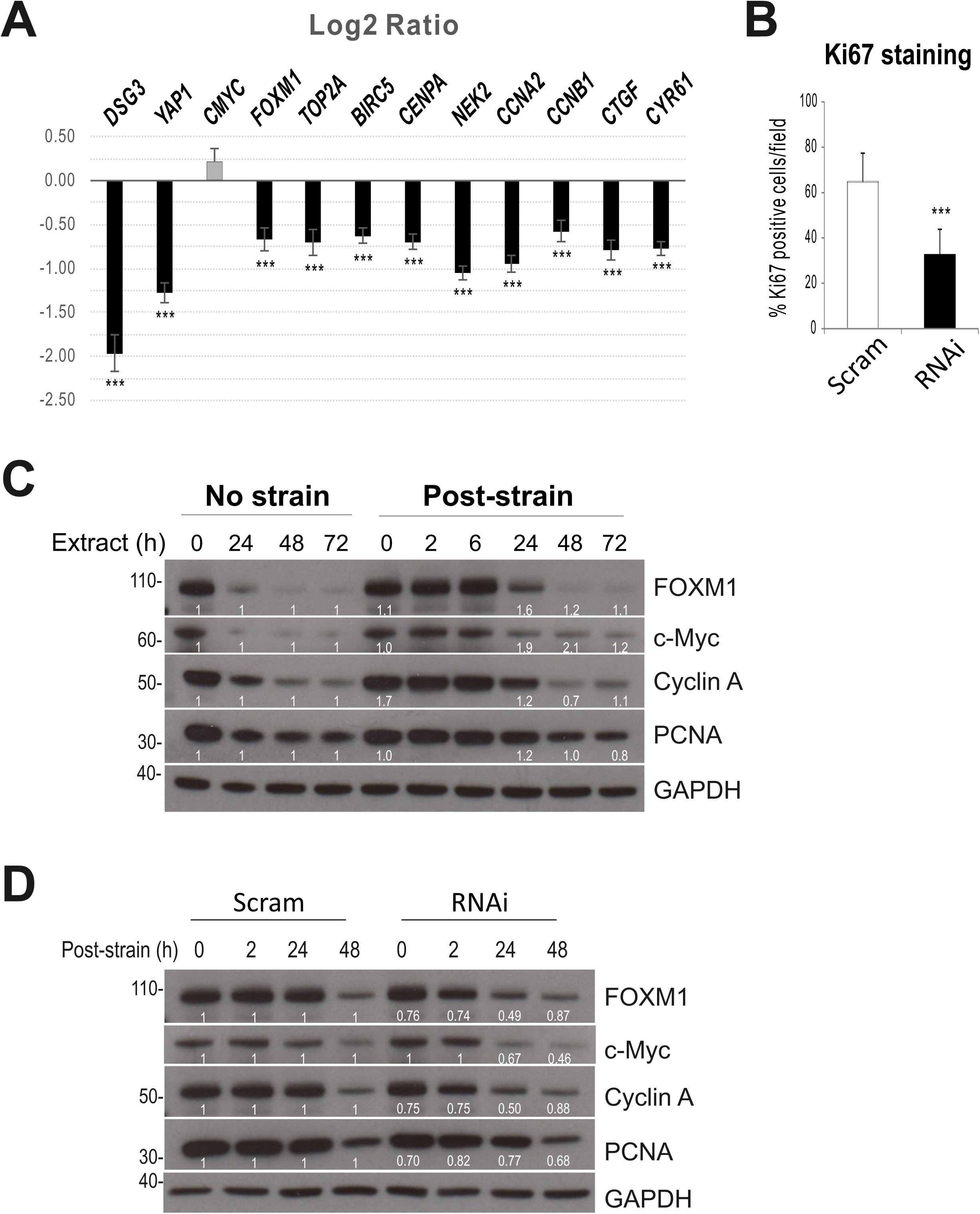
Dsg3 regulates the expression of *YAP1* and its target genes as well as other cell proliferation markers in confluent cultures. *A*, RT-qPCR analysis of cell cycle associated genes, including YAP targets such as *FOXM1, CCNA2, CENPA, CYR61* and *CTGF* in HaCaT cells with Dsg3 knockdown. Data are the log2 ratio of RNAi to Scram control (each box was the median of n=6, error bars: S.E.M., ****p<0.0001) and are representative of two independent experiments. Except for *CMYC*, all other genes showed a significant difference compared to control. Cyclic strain showed no effect on these genes in current setting, *i.e.* confluent cells subjected to cyclic strain for 6 hours before being extracted straightaway for qPCR analysis (data not shown). *B*, Ki67 staining in scrambled control siRNA (Scram) and Dsg3 siRNA (RNAi) treated cells (n=6, mean ± S.D., ***p<0.001). *C,D*, Western blotting for the indicated cell proliferation markers in obtained lysates (Fig. 3) (*C*) as well as in siRNA pre-treated cells subjected to cyclic strain for 6 hours (*D*). GAPDH was used as a loading control. The fold changes (normalized against the loading control in each lane) in post-strained cells were calculated as opposed to their respective control lanes and were displayed below of each blot.

## 4. Discussion

This study shows, for the first time, that Dsg3 cooperates with E-cadherin and other characterized mechanosensory proteins, such as α-catenin and nonmuscle actomyosin, in keratinocyte response to external mechanical forces. Importantly, our findings have identified that Dsg3 serves as an upstream regulator of YAP/pYAP and forms a complex with and sequesters pYAP to the plasma membrane, a process that involves PKPs, to facilitate AJ assembly (Fig. 10). A recent report has identified that the desmosomal plaque proteins PKP1/3 bind to 14-3-3γ/σ isoforms that are required in desmosome adhesion [42]. Another independent study has shown that PKPs interact directly with the cytoplasmic tail of Dsg1-3 [4]. This study adds these recent findings and provides direct evidence that Dsg3 is involved in complex formation with, and sequestering of, pYAP with targeting of the complex to the keratinocyte surface. This Dsg3/YAP pathway had a positive influence on the expression of YAP target genes associated with cell proliferation. Furthermore, we provide preliminary evidence that oral keratinocytes are more mechanosensitive than cells derived from the skin since many junctional proteins, including both E-cadherin and desmosomal cadherins, were stimulated by mechanical strain; reactions which showed retardation in skin keratinocytes. In summary, this study uncovers a novel signaling role for Dsg3 as a previously unsuspected key player in keratinocyte mechanosensing and mechanotransduction that potentially involves in the E-cadherin/α-catenin signaling axis.

**Fig. 10.**
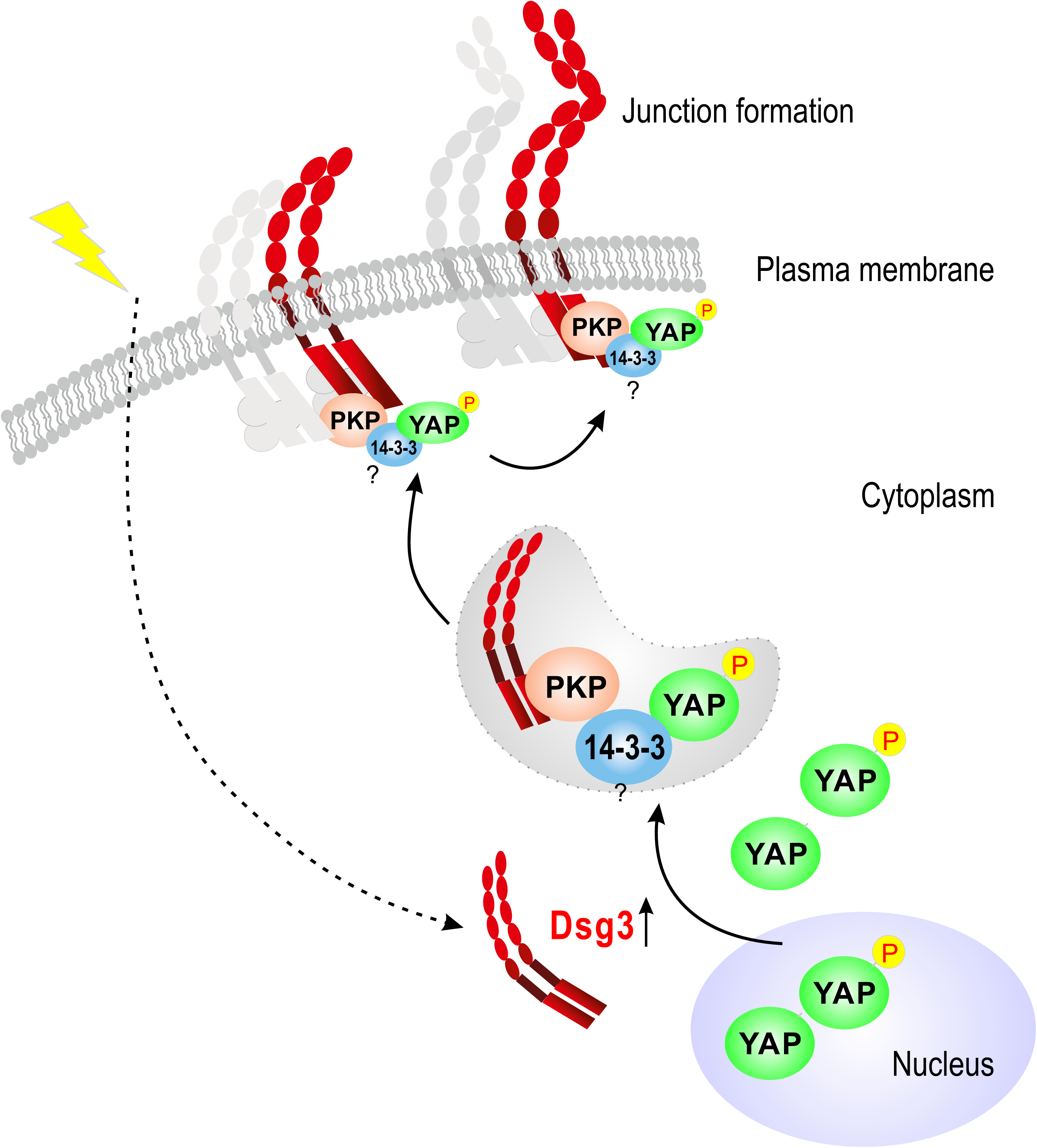
A schematic model illustrating environmental mechanical forces induce Dsg3 expression that leads to YAP/pYAP nuclear export. Dsg3 preferentially forms a complex with pYAP in the cytoplasm and sequesters it to the cell surface to facilitate junction formation in keratinocytes.

Many studies have indicated that AJs, in association with the cytoskeleton, serve as mechanosensors/mechanotransducers and are adaptable to various tension and tugging forces by modifying the cell adhesion strength, size, and protein concentration as well as causing molecular conformational changes at the junctions [12,13,26,29,30,32,57]. Emerging evidence reveals that the extra-junctional Dsg3 cross-talks with E-cadherin and regulates AJ assembly and adhesion [7,35,43,54,55]. This study demonstrated that the loss of Dsg3 action had a major impact on the junction assembly of E-cadherin and Myosin IIa in response to mechanical loading. However, rather than enhancing the cell-cell adhesion strength, we found (based on our current settings) that cyclic strain provoked junction remodeling (accelerated Dp turnover) which is consistent with previous reports that indicating negative feedback in response to large scale mechanical stresses [13,45]. Our results also showed variations in response in keratinocytes derived from different body sites. While oral keratinocytes appeared to be more responsive to mechanical stress, exhibiting an instant increase in the expression of a variety of junctional proteins, skin keratinocytes showed a delayed response during the relaxation time period with Dsg3 peak levels achieved later *i.e.* 48 hours after mechanical loading. Such a delayed response could well reflect a sustained effect after the abrogation of stretch, or as a result of “repairing” the monolayer after cyclic strain. Alternatively, this could be due to the mechanical memory of keratinocytes as reported recently [36,59]. Distinct behaviors and cellular properties between oral and skin keratinocytes have been appreciated and these include aspects of wound healing and angiogenesis [9,50,56]. Given the distinct expression patterns in Dsg3 staining between oral and skin tissues, we feel this could in part reflect their different mechanical environments.

YAP has been identified as a mechanosensor independent of the Hippo pathway, with its nuclear relocation being regulated by mechanical strain and substrate stiffness involving Rho GTPase activity and tension of the actomyosin cytoskeleton [16,24,37,39,41]. Unexpectedly, and apparently paradoxically, nuclear retention, or retardation of nuclear export, of YAP/pYAP was detected by immunofluorescence in Dsg3 knockdown cells at low cell densities on coverslips but also in confluent cultures in Flexcell wells subjected to cyclic strain. These data suggest a complex situation in which Dsg3 knockdown correlated with YAP nuclear retention but inhibition of target gene expression. We have not examined putative mechanisms but such inhibition of YAP might be caused by pronounced attenuation when expression levels fall below the threshold values [16,39] or when localization of YAP/TAZ is controlled by additional factors such as cell density and geometry [37,39]. Our previous studies have shown that Dsg3 silencing results in cell morphological changes with flattening due to defects in F-actin organization and cell polarization [35]. Thus the YAP nuclear retention in Dsg3 knockdown cells could be caused, partly at least, by cell flattening that potentiates the ECM-nuclear mechanical coupling leading to YAP nuclear location [17]. It remains unclear exactly how Dsg3 regulates YAP expression. This could be due to Dsg3’s ability to activate various signal pathways, including Src, as well as c-Jun, that regulate the actin and actomyosin cytoskeleton [7,35,43,54,55]. Dsg3 depletion has a negative impact on the formation and tension of F-actin [35,53,54] that may attenuate YAP, as supported by our qPCR data indicating reduction of *YAP1* target gene expression. In our study, however, cyclic strain did not affect the *YAP1* target gene expression nor cell proliferation for up to 3 days in confluent cultures. Nuclear pYAP-S127 location is regulated via S128 phosphorylation of YAP driven by the activation of Nemo-like kinase (NIK) [22] though precisely how Dsg3 silencing impairs epithelial cell proliferation remains unknown [34]. Probably this is due to the downregulation of YAP but this differs from α-catenin knockdown in mammalian cells which activates YAP/TAZ [27,60].

Although pYAP is regarded as an inactive protein our study suggests a potential role for pYAP in junction formation in keratinocytes in which the cell-cell junctions are particularly abundant. Various junctional proteins, such as α/β-catenins, Crumbs polarity complex, Angiomotins, and Zonula Occludens proteins, are known to physically interact with YAP/TAZ and regulate its activity [33]. This study provides the first evidence that a member of a transmembrane protein and cadherin superfamily, Dsg3, can physically interact with a protein complex containing pYAP/14-3-3/PKP1/3. As illustrated in Fig. 10, the expression of Dsg3 is stimulated by mechanical cues from cell surroundings and Dsg3 binds the pYAP protein complex in the cytoplasm and sequesters it to the plasma membrane to facilitate the junction formation. Thus we observed isochronous expression profiles of YAP, pYAP, and Dsg3 during the course of mechanically induced activation of the Hippo-pYAP pathway in the post-strained cells or the transient onset of Hippo-pYAP in cells at steady state, grown from sub-confluence to over confluence. In contrast, other junctional proteins such as E-cadherin, Dp, PKPs, and Pg did not show the same expression pattern. Dsg3 depletion caused a failure in strain-induced pYAP activation or recovery. Hippo pathway is a prerequisite for contact inhibition of cell proliferation (CIP) that controls tissue homeostasis and organ growth [2,39]. CIP is regarded as a classical paradigm of epithelial biology and its loss contributes to cancer development and progression. The previous study identifies that α-catenin is a central component in CIP and a key suppressor of YAP [27]. This study provides evidence of a novel pathway of Dsg3/YAP that differs from α-catenin in regulating the Hippo-YAP signaling. Accumulated evidence points to Dsg3 acting as a signaling molecule and this study connects its signaling role to control keratinocyte proliferation, and therefore organ size, via the mechanism of regulating YAP [16,39].

Notably, two independent reports recently demonstrated that the desmosomal proteins, Dsg2 and Dp function as the biosensor in human cardiomyocytes and canine kidney simple epithelial cells [3,40]. These findings, along with the current study, collectively underscore the importance of desmosomes as mechanosensory and load-bearing structures that coordinate with AJs in control of tissue integrity and homeostasis.

## Acknowledgments

We are very grateful to Ian R. Hart for his assistance with editing of the manuscript. We thank Steven Thorpe and Diana Blaydon for assistance in manipulating the Flexcell Tension system, Ken Parkinson and Matthew Caley for the antibody reagents, Catherine Harwood for providing T8 cell line, Belen Martin-Martin for assistance with super-resolution microscopy, Fiona Kenny and William Ogunkolade for technical help, and Simon Rawlinson for critical reading of the MS. This work was supported by a scholarship from Naresuan University, Phitsanulok, Thailand.

## Authors contributions

H.W., J.U., U.S.A., A.R. and Y.H. investigation and validation; H.W., J.U. data curation; J.U., H.W., U.S.A. and M.T.T. formal analysis, J.U. funding acquisition; H.W. project administration and supervision; J.T.C. and A.C.L. supervised and assisted with the PA gel study, M.T.T. assisted with the qPCR analysis, A.K. and R.C. helped with the plasmid amplification, J.S. helped with confocal image analysis; J.T.C. and M.T.T. discussed the obtained data and helped with the manuscript construction, H.W. and J.U. wrote, review and editing manuscript.

